# Behavioral and neural correlates of social hierarchy formation in a sex-changing fish

**DOI:** 10.1101/2024.07.04.602141

**Authors:** Haylee M. Quertermous, Kaj Kamstra, Chloé A. van der Burg, Simon Muncaster, Erica V. Todd, Christine L. Jasoni, Culum Brown, Neil J. Gemmell

## Abstract

Social hierarchies in sex-changing fish determine which fish will change sex, yet the complexities of hierarchy formation at the neurobehavioral level are still being unraveled. Here, we investigate the formation of social hierarchies within groups of New Zealand spotty wrasse, integrating behavioral observations with neural activation patterns upon social disruption. We find that dominance hierarchies form linearly based on size, with larger fish displaying more dominant behaviors and smaller fish displaying more submissive behaviors. Disruption of the social hierarchy induced rapid behavioral changes, particularly in second-ranked fish, highlighting that second-ranked fish will opportunistically adopt a dominant position. Analysis of neural activation patterns reveals that the social decision-making network is deeply involved in the establishment of dominance, with the fish attaining dominance showing significant differences to all other ranked fish. Overall, this study underscores the complexity of social relationships and their neural underpinnings in the spotty wrasse, providing a foundation for further research into the cellular and molecular mechanisms of socially-controlled sex change, and demonstrates that disruption of the social hierarchy triggers rapid changes in both behavior and the social decision-making regions of the brain.

## Introduction

Social dominance hierarchies are common across the animal kingdom and have important implications for survival and reproduction. Dominance is associated with asymmetric aggression and priority access to resources that ultimately increase fitness (e.g., mating opportunities, food, shelter) (1). One of the most extreme consequences of a dominance hierarchy is socially-controlled sex change. In protogynous (female-to-male) species, removal of the dominant male will trigger sex change in the dominant female. These fish are often polygynous (2,3), with a single male maintaining a territory or breeding ground with multiple females (4). While it is the male that holds the most dominant position, the females can arrange themselves further in the hierarchy (5–7). Often, the dominant female will change behavior within minutes to hours after removal of the male, with gonadal and morphological changes following on a scale of days to months (8,9). The main hypothesis as to why sex change occurs is outlined in the size-advantage model, suggesting body size influences reproductive success differently between sexes, and that an individual can maximize its reproductive success by changing sex once a particular body size is reached (10,11). Therefore, individuals should first reproduce as a certain sex (e.g., female) early on and change sex when they have increased to a size or age that they will benefit as the opposite sex (e.g., male). Thus, size plays an important role in hierarchy formation as has been shown in other fishes (12,13), yet this has scarcely been systematically tested in sex-changing fish (14–16).

Although changes in social environment are known to be the primary triggers for sex change, most research on sex-changing fish considers only behavior among the most dominant individuals, with little attention for the entire group (17). Understanding differences between subordinates could provide more insight into earlier changes that occur before sex change. It remains unknown if there are factors that delineate high-ranked individuals predisposed to change sex from conspecifics, and whether these differences prime the molecular and neural systems that underlie sex change. If the second-ranked individual can ‘determine’ that they will be the most fit for sex change even before male removal, this may initiate molecular changes priming them for sex change. Although the costs and benefits of being dominant and subordinate are well-studied (18–22), more work is needed to observe differences between subordinates of different rank (23). Understanding the behavioral differences between females of different ranks among protogynous hermaphrodites will help us determine how female dominance hierarchies are formed, maintained, and altered. While there is some past research on female hierarchy formation in fishes, this is dominated by work on cichlids (24,25), and our understanding of how female hierarchies form and their role in sex-change in sequential hermaphroditic fishes is lacking.

It is clear that socially-controlled sex change critically relies on changes and signaling in the brain, independent of the gonad; in bluehead wrasse, ovariectomized fish still change sex, both behaviorally and externally (26). The complex network of neuroendocrine pathways has also been well explored, including the role of steroid hormones, triggering of the HPI and HPG axes, and the role of neuropeptides (e.g., AVT, IST) (27). The preoptic area (POA) in the hypothalamus has also been shown to have a key role in sex differentiation and sex change in anemonefish (28). Recent work has shown that new cells are added to the POA during protandrous sex change in the anemonefish *Amphiprion ocellaris* (29). Further, courtship behavior in protogynous bluehead wrasse has been well studied, showing to be controlled by a network of brain areas associated with the social behavior network: a highly conserved network of brain regions suggested to control social behaviors such as reproduction, aggression, and parental care. (30,31). While many of these studies have provided evidence for the neuroendocrinological mechanisms underpinning sex change on a broader temporal scale, a brain-wide investigation of early neural activity upon disruption of an established and stable social hierarchy is lacking. Neural activity of hierarchy formation and social dominance has broadly been studied in males (22,32–34), but female hierarchies have largely been neglected (35).

We investigate here the group formation of New Zealand spotty wrasse (*Notolabrus celidotus*; i.e., “spotty”), a protogynous sequential hermaphrodite. Spotty are ubiquitous across New Zealand and amenable to captivity (36,37). Spotty have three sexual phenotypic morphs: initial phase (IP) females, IP males, and terminal phase (TP) males. TP males and IP individuals can be easily differentiated due to their different color morphs. All spotty first have IP coloration, with most juveniles developing as females and a small minority as IP males (37). During sex or role change to a TP male, individuals change coloration. TP males are typically in groups with 4-6 females (37). We allocated spotty into novel groups of 5 females and determined hierarchical rank based on submissive and aggressive behavior. Subordinate and dominant behaviors were analyzed over nine days. After 11 days, we removed the most dominant fish from each group and analyzed the behavior of the remaining fish for 30 minutes. One hour after social disruption, we removed and sampled fish to measure neural activation, using pS6 immunohistochemistry in the brain, based on the observed rank and behavior. We predicted that dominance and size would correlate linearly (i.e., the bigger the fish the more dominant) and that the more dominant fish would show increased neural activation in brain regions controlling aggression in comparison to subordinate fish.

## Materials & Methods

### Animals

Animal handling was performed in accordance with New Zealand National Animal Ethics Advisory Committee guidelines. Spotty were captured around high tide by hook and line off the coast of Tauranga, Bay of Plenty, New Zealand (37.6878° S, 176.1651° E) and subsequently maintained at the Aquaculture Centre at Te Pūkenga/Toi Ohomai Institute of Technology, Tauranga. Fish were housed for several months in large 1600L recirculating holding tanks to acclimate prior to experimentation. Prior to experimental tank allocation, standard length (SL; ranging from 108.87-179.91mm, Figure S1) and body weight (ranging from 22-122g) were measured. SL was measured by taking photos of each fish next to a metric ruler and using Image J; two measurements from the tip of the nose of the posterior margin of the caudal peduncle were taken in Image J and then averaged (Table S1). Floy tags (Fine Anchor T-bar 5 cm, Hallprint T-Bar tags) were attached to enable individual recognition. Fish were housed in 400-L recirculating seawater systems under a natural photoperiod. Fish were fed frozen greenshell mussels (*Perna canaliculus*) two to three times per week. At the conclusion of behavioral experiments, fish were anaesthetized through immersion in 600 ppm 2-phenoxyethanol and euthanized by decapitation.

### Experimental Design

#### 1. Group Formation

Eleven groups of five IP fish of different sizes were placed in 400-L recirculating seawater systems and kept under a natural photoperiod. A PVC pipe (∼100mm wide x ∼200mm long) was placed in each tank to act as a shelter. Behavior was recorded 2-3 times per day between 09:00-18:00 for nine days after group initiation. Videos were taken using GoPro HERO Session and GoPro Hero 11 Black Mini cameras for a period of 20-30 minutes. Dummy cameras were placed in the tanks when recordings were not being taken to habituate the fish to the presence of a camera.

#### 2. Social Disruption

After 11-12 days of group formation, the dominant fish was removed in n=7 treatment tanks. In the remaining n=4 control tanks, the dominant fish was removed and immediately reintroduced to act as a control for the disruption. All tanks (control and treatment) were then recorded for one hour post disruption and then all remaining fish were removed and immediately euthanized for gonadal histology and immunohistochemistry analysis. See Figure S2 for a diagrammatic description of this process.

### Behavioral Analysis

Videos from group formation and social disruption were watched and scored using BORIS software (38). For group formation, all videos during the first three days, and then videos from every other day (days 5, 7, and 9) were analyzed. All individual observations were summed together for statistical analysis. For social disruption, the first 30 mins of recording was analyzed. At the start of all recordings, 10 minutes were left unanalyzed to allow for fish to recover after the introduction of the camera. The next 10 minutes were then observed to count the number of aggressive and submissive interactions. A rush (aggression) was defined as a directed movement or acceleration towards another individual and an escape (submission) as acceleration away from or change in direction to avoid another individual. Position and movement in the tank were also recorded. State behaviors were measured in seconds and were divided into three behavior types: Time spent in shelter, time spent on the floor, and time spent swimming around the tank (Table S2b).

Each individual was given a dominance score (DS) calculated by their overall rate of rushing behavior divided by their overall rate of rushing plus the overall rate of escaping behavior (rRush/(rRush+rEscape)). Therefore, highly dominant individuals will have a DS closer to 1 and highly subordinate individuals a score closer to 0. This dominance index is similar to one used by Oliveira and Almada (39), but uses the rate of aggressive and submissive interactions rather than the number of behaviors. The proportions and rates of the behaviors were used, rather than the measure of time or total number of a point behavior by itself, as some fish were out of view for periods of filming, therefore, not all fish have the same total time of observation. Fish in each tank were given a rank from 1-5 (1 being the most dominant) based on their DS.

### Gonadal histology

Gonadal samples collected upon finalization of the social disruption experiment (see subheading “Social Disruption”), were fixed in 4% paraformaldehyde in 1× PBS solution at 4°C for 72 hours. Gonadal samples were embedded in paraffin and subsequently cut at 3–4 μm on a Leica RM2125 RTS rotary microtome. Standard Mayer’s haematoxylin and eosin staining was performed as previously described (36) to assess tissue structure. Images were taken using an Aperio CS2 Digital Slide Scanning System (Otago University Histology Unit). Gonadal sex was classified according to Goikoetxea et al. (40).

### Immunohistochemistry

Following social disruption, brains from both control and treatment rank 2-5 fish were dissected and subsequently fixed in 4% paraformaldehyde in 1x PBS solution at 4°C for 72 hours, then brains were cryoprotected in 30% sucrose solution until the tissue lost buoyancy. Brains were flash frozen in dry ice-cooled isopentane and stored at -80°C. Brains were embedded in optimal cutting temperature medium (TissueTek, Sakura, Torrance, CA, USA) and sectioned coronally at 50μm on a CM1950 Leica cryostat. Cryosections were collected on alternate sets of SuperFrost plus slides (Thermo Fisher Scientific NZ Ltd, North Shore City, New Zealand), air dried at room temperature overnight and stored at -80°C. Slides were thawed and cryosections were surrounded with a hydrophobic barrier. Slides were then washed 3 times for 10 minutes with 1× PBS. No endogenous peroxidase activity was detected following a pilot incubation with the chromogenic substrate solution. Non-specific binding was blocked (1% bovine serum albumin, 0.3% Triton-X and 5.0% normal goat serum made in 1× PBS) for 2 hours at room temperature, and slides were incubated with primary pS6 antibody (1:1500; Cell Signaling Technologies pS6 ribosomal protein S235/236 rabbit monoclonal antibody #4858) overnight at 4°C. Phosphorylated S6 protein (pS6) can be employed as an indicator of neural activity (41), as it reveals cytoplasmic staining of ribosomal proteins that were phosphorylated in the preceding hour, a process associated with enhanced translation within the cell. The pS6 marker has been utilized in various fish species across different social or sensory contexts in the brain (42,43). Slides were rinsed with 1× PBS (3×10min), and further processed using the Vectastain ABC-HRP Kit (PK-4001)) according to the manufacturer’s instructions. The secondary antibody used was a biotinylated goat anti-rabbit IgG (1:200, PK-4001, Vector Laboratories). To visualize the staining, a HRP peroxidase substrate kit (Diagnostic BioSystems #K062-110) was used according to the manufacturer’s instructions. Finally, slides were dehydrated through an alcohol series (50%, 70% and 95% for 1 min each; 100% 2×2 min each), cleared with xylene (2×2 min) and coverslipped using dibutylphthalate polystyrene xylene. Specificity of the staining was confirmed through omission of either the primary antibody (Figure. S3B) or the secondary antibody (Figure S3C) and through the use of a pS6 blocking peptide (Cell Signalling #1220; Figure S3D). Nissl counterstaining confirmed cytoplasmic labelling (Figure S3E). Detailed counterstaining for each ROI shown in Figure S4.

### Brain image analysis

Photomicrographs were taken on a Nikon Ti2-E inverted bright field microscope with a 20x magnification objective. Composite images were taken with 15% overlap and stitched together using Nikon NIS-Elements software. Raw images were processed in Fiji/ImageJ and Adobe Photoshop (Adobe Systems Inc., San Jose, CA) to correct background noise and minor tissue artefacts. Image segmentation was performed as described previously using the Trainable Weka Segmentation plugin (44). Following comparison of the images with the spotty wrasse brain atlas (45), cell counts were performed using the Automatic Cell Counting with Trainable Weka Segmentation tool (46). The density of pS6-stained cells was calculated as the number of stained cells divided by the surface area of the regions of interest quantified. For each region (Table 17c), 3–4 consecutive sections were quantified (dependent on region, but consistent across animals) at the same rostro-caudal location within the nucleus across animals and averaged together for each animal to obtain a mean pS6-stained cell density per brain region (# of cells per μm^2^). Brain areas that were identified to have significantly different pS6 cell counts between ranks were recounted manually by an experimenter blind to the experimental parameters. Several of the brain regions implicated in the current study contain a multitude of different cell types and consist of multiple subregions. Because the pS6 staining does not differentiate between cell types we did not analyze these subregions separately.

### Statistical analysis

Behavioral data (group formation and social disruption) was analyzed in R 4.1.1, using the R Studio Interface Version 2023.12.1+402. The alpha level was set to 0.05; however, a p-value of 0.1 was noted as marginally significant. Linear mixed-effects models (LMMs) were used to determine how behavior variables were associated with DS, average SL, and the interaction between DS and SL. Tank was set as a random effect for all models. We then used models with rank as the fixed effect to get contrasts between all ranks. The *lmer* function was utilized from the *lme4* package. Normality was tested using the Shapiro-Wilk test. The proportion of time on floor (pFloor), rate of rushes per minute (rRush), and rate of escapes per minute (rEscape) variables were square root transformed to obtain a normal distribution. The best fit models (i.e., lowest AIC) were determined using the *aictab* function in the *AICcmodavg* package (Tables S4-8). Immunohistochemistry data analysis was performed using Prism (GraphPad Software, La Jolla, CA). Comparison of cell counts between brain areas was done by one-way ANOVA, followed by Dunnett’s multiple comparison test, comparing the “rank 2” group to all other groups. Results are presented as the mean ± SEM. *p* < .05 was considered statistically significant.

### Brain-behavior correlation analysis

Behavioral and neural data obtained during social disruption were analyzed together to identify which regions of the brain correlate with which behaviors, by social rank. Behaviors recorded during group formation were not used for these analyses. Outlier and/or confounding behaviors (see Supplementary Materials notes) were first removed from analyses, and fish were only used if complete neural and behavior data was available (see Table S11) for a total of 37 fish (four control tanks and six treatment tanks). No rank 1 fish (most dominant fish) were used in these analyses, as brain regions were not analyzed in these fish. The data were analyzed in R (v 4.3.1), using Rstudio (v 2023.12.1+402). PCA plots were generated for behavior and neural data separately, using the *prcomp* function with scale=TRUE and plotted using *ggplot2 v3*.*5*.*1*. The loadings from principal component 1 and 2 were extracted using the *hi_loadings* function in pcaExplorer (v 2.28.0). Scree plots were generated by plotting all principal components against extracted eigenvalues (prcomp_object$sdev^2), using *ggplot2*. Scree plots were then used to determine which PCs had an eigenvalue > 1 and all combinations of behavioral PCs and neural PCs were then compared, with statistical analysis performed using Pearson’s product-moment correlation. Sample correlation plots were generated using the *heatmaply_cor* function with na.rm=FALSE in heatmaply (v 1.5.0). Statistical differences between ranks (control versus treatment treated separately) were calculated using the *pairwiseAdonis* wrapper (v 0.4.1) with 999 permutations. As PCA showed that ‘rank 2 treatment’ samples form a clear separate cluster, statistical differences between the ‘rank 2 treatment’ samples and all other samples were also calculated using *adonis2* function in vegan (v 2.6.4) with method=‘eu’.

## Results

### Dominance hierarchy in the New Zealand spotty wrasse is rapidly established based on size

A piecewise linear model was utilized for the relationship between DS and size (SL) with a breakpoint at 140mm (Figure S5). SL was significantly and positively associated with DS, with larger fish having higher DS, both for fish above (β = 4.13e-3, df = 1,45.82, t = 2.02, p = 0.050) and below 140mm (β = 0.02, df = 1,41.58, t = 16.36, p < 0.001). The SLs were significantly different between all ranks, except for 3 and 4, with fish being significantly longer than fish ranked subordinate to them (Figure S6; Table S9). As expected, with rank being given based on DS, all ranks had significantly different DS from one another (Figure 1C; Table S10). A clear dominance hierarchy was apparent almost immediately, with only ranks 3 and 4 not being obvious in the first observation, but quickly differentiating after just a few observations (Figure S7). All raw group formation behavioral data can be found in Table S2.

**Figure 1.**
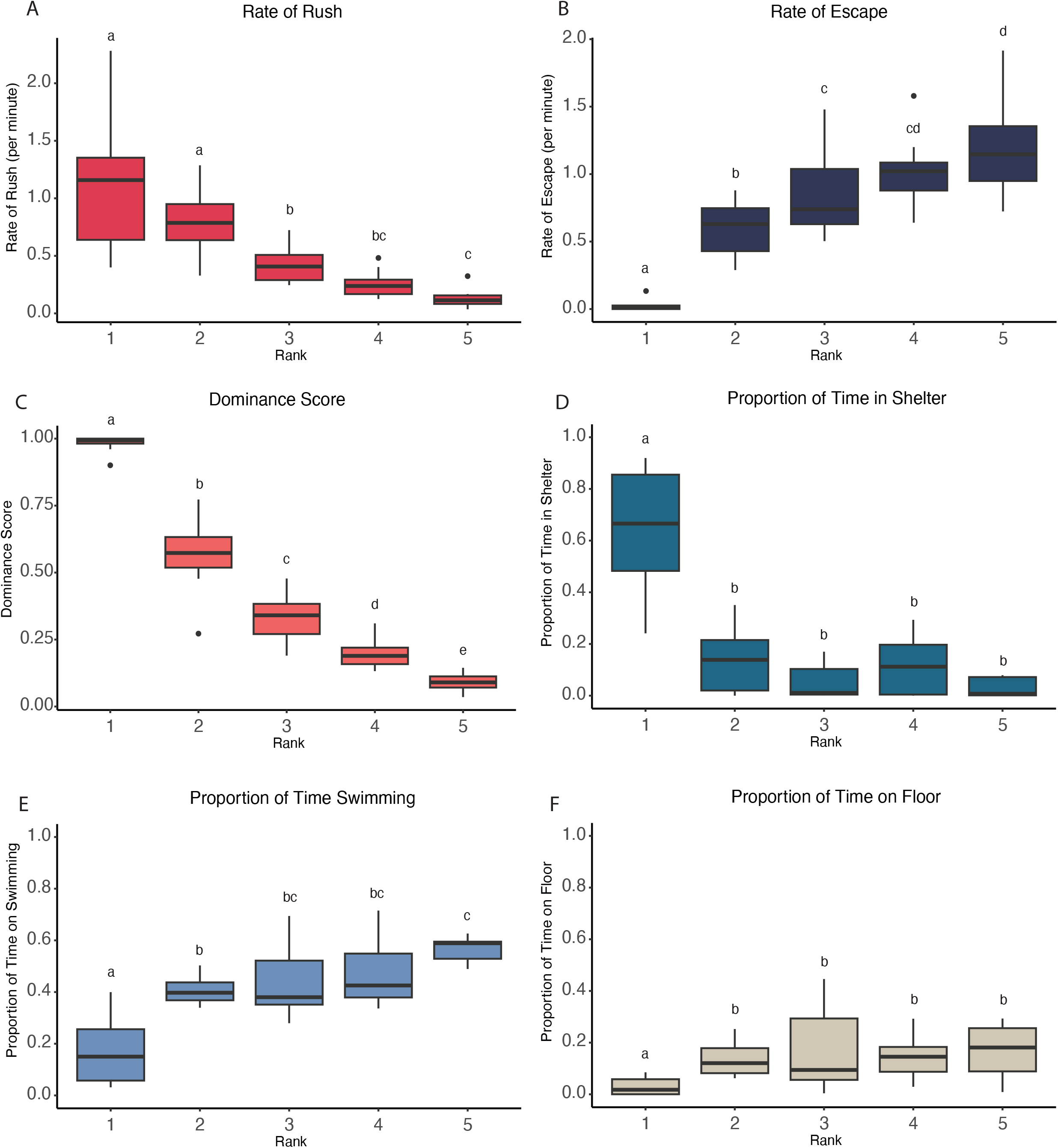
Group formation behaviors by social rank. (A) The rate of aggression (i.e., rushing behavior) decreases moving down the hierarchy. (B) Submission (i.e., escaping behavior) increased moving down the hierarchy. (C) Dominance score is significantly different between all fish ranks. (D) Rank 1 spent significantly more time in the shelter than all other ranks and significantly less time swimming (E) and on the floor (F).

### Group formation behaviors are dictated by dominance hierarchy

DS was significantly and positively correlated with proportion of time spent in the shelter (pInShelter: β = 0.60, df = 1,53, t = 7.73, p < 0.001). However, when rank replaced DS as the fixed effect (Table S4), the model had a better fit and demonstrated that the fish with the highest DS (i.e., rank 1) spent significantly more time in the shelter than all other ranks (Table S11). The other ranks did not differ significantly from one another (Table S11). The best fit LMM for proportion of time spent swimming (pSwimming) included only DS as a fixed effect (Table S5). Fish with a higher DS spent a significantly smaller proportion of their time swimming (β = -0.37, df, 1,53, t = -7.62, p <0.001). The model with rank as the fixed effect did not have a better fit and showed that rank 1 spent significantly less time swimming than all other ranks and that rank 2 spent significantly less time swimming than Rank 5. The rest of the ranks did not differ significantly from each other (Table S12). The best fit LMM model for the proportion of time on floor (pFloor) also only had DS as a predictor variable (Table S6). DS and pFloor were significantly and negatively correlated with each other (β = -0.25, df, 1,43.39, t = − 4.83, p <0.001). Using rank as the only fixed effect did not have a better fit model, but it showed that Rank 1 spent significantly less time on the floor than all other ranks, and that all the other ranks did not significantly differ (Table S13). Figure 1D-F illustrates the proportion of time each rank spent on each of the three major activities (pInShelter, pSwimming, and pFloor), which shows that fish of decreasing rank spend more time swimming, and rank 1 fish spend the most time in the shelter. Using rank as a fixed effect in the model testing the rate of rushes per minute (rRush), we show that rank 1 had a significantly higher rRush than ranks 3-5, but did not differ significantly from rank 2. rank 2 also had a significantly higher rRush than ranks 3-5. Rank 3 had a significantly higher rRush than rank 5, but did not differ significantly from rank 4. Rank 4 and 5 did not differ significantly from each other (Table S14). For rate of escapes per minutes (rEscape) between ranks, which was the best predictor variable, we show that rank 1 had a significantly lower rEscape than all ranks and rank 2 also had a significantly lower rEscape than ranks 3-5. Rank 3 had a significantly lower rEscape than rank 5, but did not differ significantly from rank 4. Rank 4 and 5 did not differ significantly from each other for rEscape (Table S15). Figures 1A and 1B illustrate the rates of rush and escape behaviors across each rank, which clearly shows increasing submissive (escape) behavior as the ranks decrease, and inversely, decreasing aggressive (rush) behavior as the ranks decrease. All model AIC data for rRush and rEscape are in Tables S7-S8.

### Removal of the dominant fish induces a linear shift in hierarchy status

In treatment tanks, after the removal of the rank 1 fish, rank 2 experienced a significant increase in DS and rRush, with a significant decrease in rEscape (Figure 2). Rank 3 had a significant increase in rRush but no significant differences for DS and rEscape. Rank 4 experienced both a significant increase in rRush and rEscape, but no significant change in DS. Rank 5 had a significant decrease in DS, a marginally significant decrease in rRush, and a significant increase in rEscape. Rank 2, 3, and 4 had a significant decrease in pFloor, and rank 5 a marginally significant decrease. Rank 3 and 5 also had a marginally significant increase in pSwimming (Table S16). Thus, fish retain their relative position in the social hierarchy, and all move up one spot.

**Figure 2.**
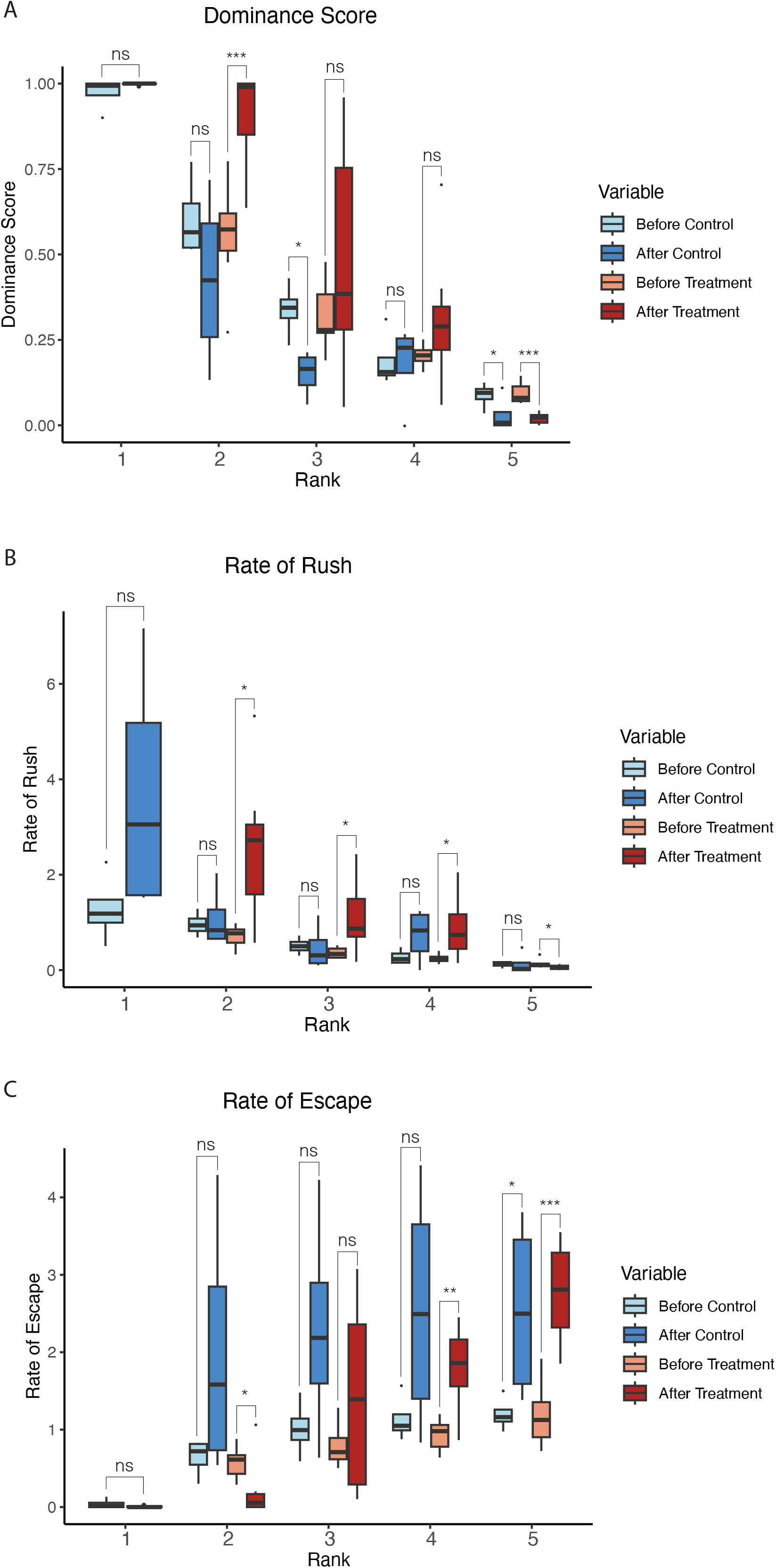
Behavior during group formation and after social disruption. Control tanks are represented in red/pink with behavior during group formation (Before Control) in pink, and after social disruption (After Control) in dark red. Treatment tanks are in blue, with light blue showing behavior during group formation (Before Treatment) and royal blue showing behavior after social disruption (After Treatment). Rank 1 does not have data for the experimental treatment as it was removed from the tank.

In control tanks, after the rank 1 fish was removed and immediately returned, rank 1, 2, and 4 did not have a significant change in DS; however, rank 3 had a significant decrease and rank 5 a marginally significant decrease in DS (Figure 2A). No rank had a significant change in rRush (Figure 2B). Rank 5 had a marginally significant increase in rEscape, while no other ranks differed significantly (Figure 2C). Rank 1 had a significant decrease in pInShelter and a marginally significant increase in pSwimming (Table S16). All other ranks did not have significant changes in any of their state behaviors. All raw social disruption behavioral data can be found in Table S3.

### Dominance is confirmed by masculinization of the gonad in most, but not in all dominant fish

In 7 out of 11 tanks, the largest and most dominant fish showed clear signs of gonadal masculinization (Figure S8), despite external coloration and morphology indicating the fish were IP females. However, given that these fish were already the largest, it is most likely that they had begun the sex changing process before the start of the experiment. Due to the identical external morphology of IP female and IP males, it is impossible to tell the two apart until gonadal morphology can be assessed. However, it is statistically unlikely that the gonad masculinization in the largest fish was because they were IP males (see Supplementary Material notes). Tank 8 was excluded from further analyses due to the presence of an IP male (see Supplementary Material notes)

### The social decision-making network is differentially activated in dominant vs subordinate fish

Neural activation patterns were compared between differently ranked fish following removal of the dominant fish. Brain areas that were identified as having significantly different pS6 cell counts between the new dominant (= rank 2) vs subordinate fish (= rank 3-5) were: the medial part of the dorsal telencephalon (Dm; Figure 3A), the dorsal part of the ventral telencephalon (Vd; Figure 3B), the ventral part of the ventral telencephalon (Vv; Figure 3C), the supracommissural nucleus of the ventral telencephalon (Vs; Figure 3D), the central nucleus of the telencephalic are (Vc; Figure 3E), the preoptic area (POA; Figure 3F), the ventral tuberal region of the hypothalamus (vTn; Figure 3G), the periaqueductal gray (PAG; Figure 3H), the periventricular nucleus of the posterior tuberculum (TPp; Figure 3I) and the anterior tuberal nucleus (aTn; Figure 3J). Putative mammalian homologs of these brain areas are listed in Table 2. In each of these brain regions the number of pS6-stained cells (Table S17a) was significantly different between the now largest rank 2 fish and any of the other fish ranked 3-5 (one-way ANOVA with Dunnett’s multiple comparisons test, Table S17b). No significant differences were observed between the fish ranked 3-5. Importantly, this difference is a result of the removal of the rank 1 fish specifically, as there were no significant differences between any of the ranked fish in the control group where the rank 1 fish was caught but immediately returned, or between the fish ranked 3-5 in the experimental group and any of the fish in the control group. In 9 out of 10 of the identified brain areas we observed increased neural activity in the rank 2 fish compared to the other fish. In the aTn, on the other hand, a decrease in pS6 staining was observed in the rank 2 fish compared to the other ranks.

**Figure 3.**
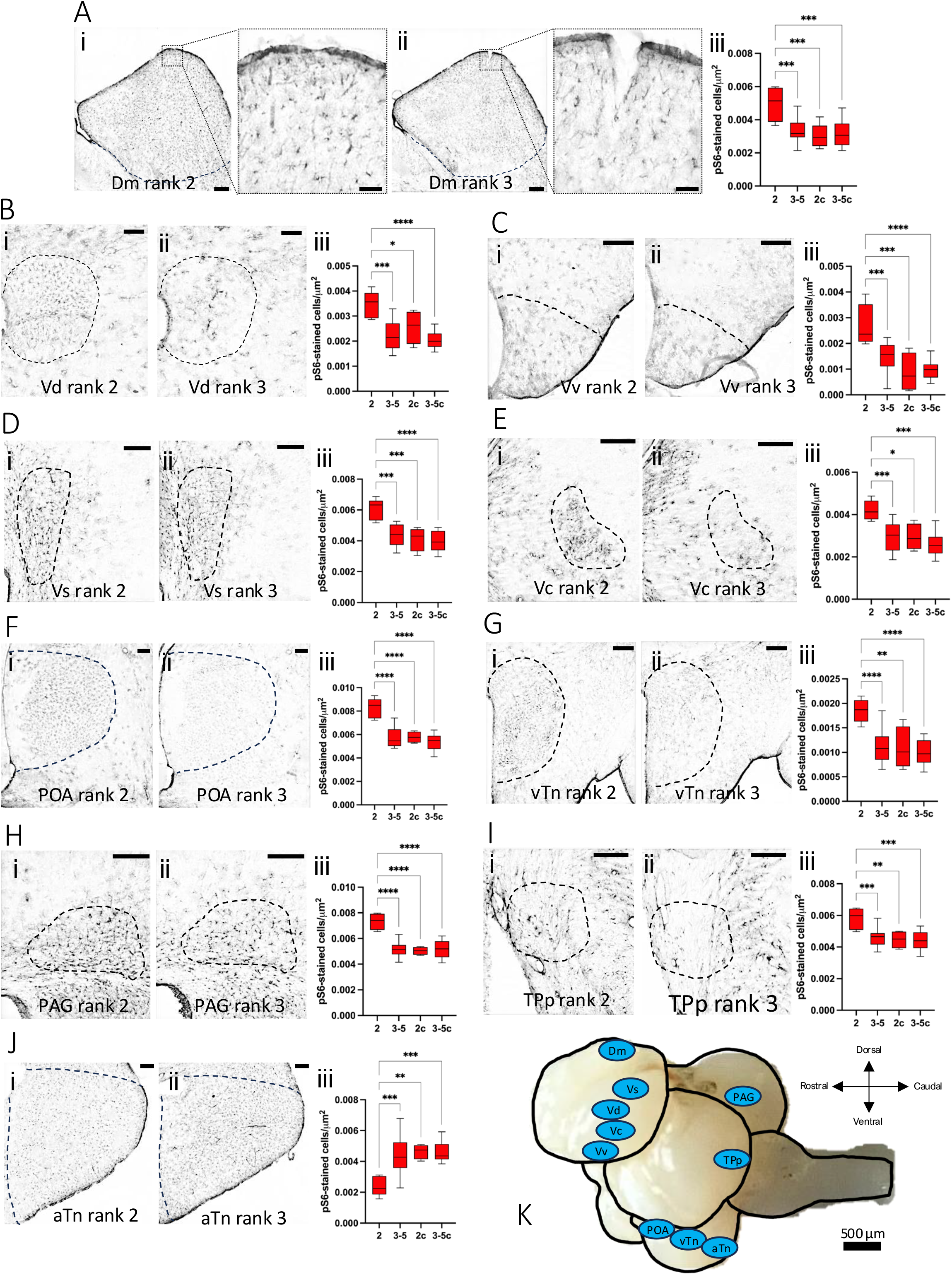
Neural activation patterns after social disruption. Representative images for each brain area are depicted on the left (I; rank 2) and middle (II; rank 3-5) in each panel. Boxplots of pS6 counts are depicted on the right of each panel, with 2c and 3-5c representing the controls (III). (A) The medial part of the dorsal telencephalon (Dm), with magnification of the insets to the right of each panel (scale bar of insets = 20μm). (B) The dorsal part of the ventral telencephalon (Vd). (C) The ventral part of the ventral telencephalon (Vv). (D) The supracommissural nucleus of the ventral telencephalon (Vs). (E) The central nucleus of the telencephalic area (Vc). (F) the preoptic area (POA). (G) the ventral tuberal region of the hypothalamus (vTn). (H) the periaqueductal gray (PAG). (I) the periventricular nucleus of the posterior tuberculum (TPp). (J) The anterior tuberal nucleus (aTn; Figure 4J). Scale bar = 100μm. (K) Overview of brain regions with significantly different pS6 counts in rank 2 fish compared to other ranks.

### Behavior and brain activity correlate based on social rank and the social decision-making network

Principal component analysis of all behavioral observations and neural activity for each of the 37 individual fish (Table S18) revealed that samples are mainly organized by social rank when considering all behavior factors (PC1: 41.2%; Figure 4A), with escape and rush behavior having the largest effect on PC1 direction (Figure 4B.i). PC2 explains only 24.8% of the variance, but a larger variation between rank 2 fish in particular is observed across this PC, which may be explained by differences in time spent rushing, swimming, and time on floor (Figure 4B.ii). Within-rank variation appears to increase with a rise in ranking (Figure 4A), reflecting the increased social instability or opportunities associated with higher ranking. Principal component analysis of neural factors (Figure 4C) shows a less distinct rank-based organization, with the primary difference in variation across PC1 (54%) being the rank 2 treatment fish forming a cluster, which appears to be driven fairly evenly by all (except one, aTn) brain regions (Figure 4D.i). Rank 2 and 3 fish from tank 6, however, appear to be ‘swapped’, in that the rank 3 fish clusters with all other rank 2 fish. This is likely due to natural biological variation and indicates that in this tank the rank 3 fish may have been more opportunistically dominant after social disruption. In fact, while fish ranks are shown here by their group formation rank (i.e., rank scored based on cumulative behaviors observed during group formation), when looking at the rank as scored only by behaviors observed during social disruption, the rank 3 fish scored higher than Rank 2 in tank 6 (see Table S3, GFRank and SDRank). Further analysis of PCs showed that for behavior, PC1-PC3 have eigenvalues > 1 and for brain, PC1 and PC2 have eigenvalues > 1 (Figure S9). All possible comparisons were performed (Figure S10), which showed only behavior PC1 vs neural PC1 to have a statistically significant (p = 9.61x10^-6^) linear relationship (r=0.66), shown in Figure 4E.

**Figure 4.**
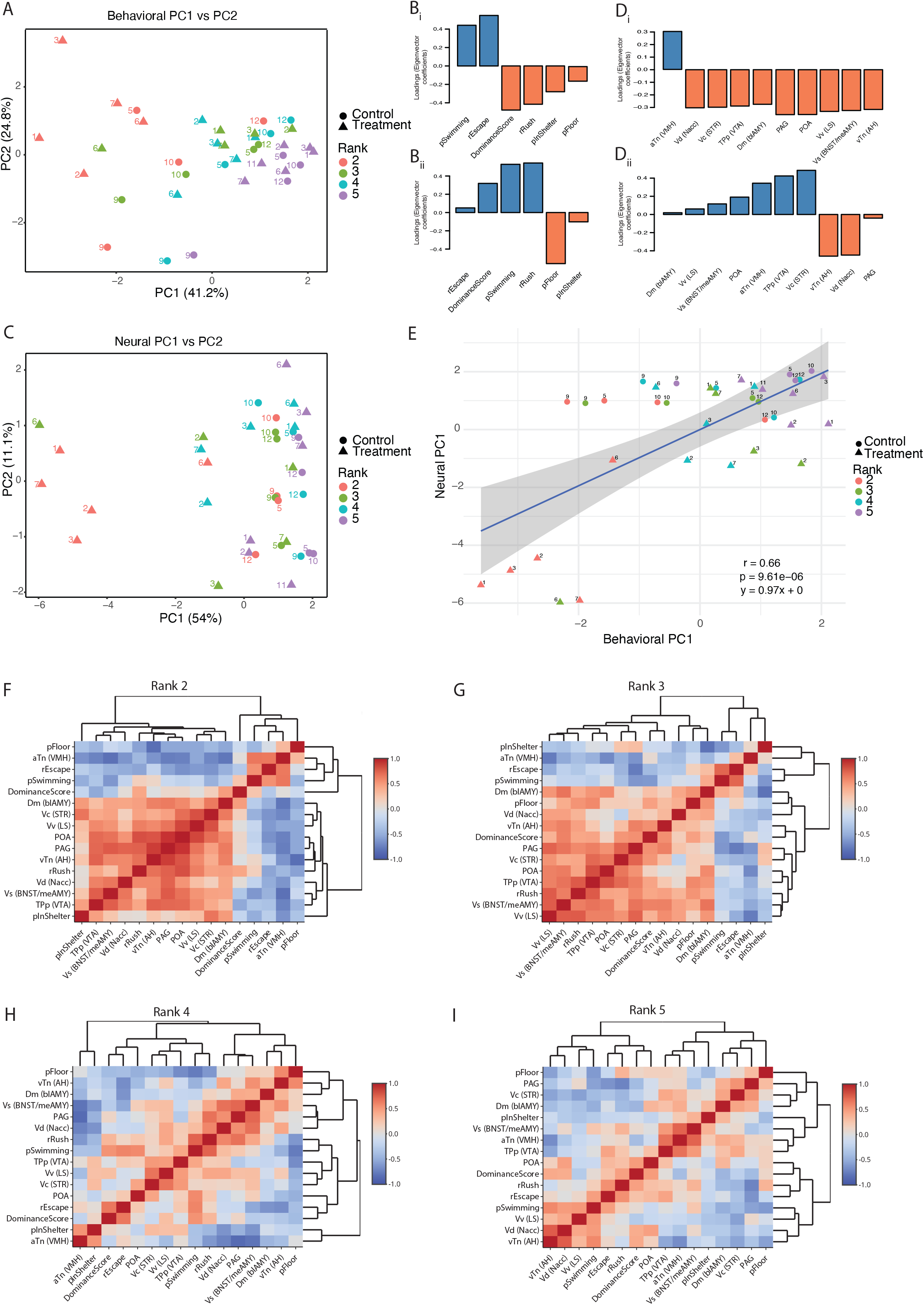
Brain-behavior correlation analyses after social disruption. (A) PCA plot of behavioral data. Sample labels correspond to tank number. (B) Loadings for each behavioral factor for PC1 (Bi) and PC2 (Bii). (C) PCA plot of neural data. Sample labels correspond to tank number. (D) Loadings for each neural factor for PC1 (Di) and PC2 (Dii). (E) Correlation of behavioral PC1 and neural PC1. (F-G) Sample correlation matrices of all neural and behavioral data, organized by rank. All correlation values are in Table S19.

Statistical analysis using all behavior and neural activity data showed that: comparing treatment samples between ranks shows only rank 2 treatment vs rank 5 treatment are significantly different (Pr > F = 0.002; Table S20). Comparing control samples between ranks shows only rank 3 control vs rank 4 control are significantly different (Pr > F = 0.001; Table S21). Comparing treatment versus control samples are only significantly different within rank 2 (Pr > F = 0.008; Table S22). However, the rank 2 treatment group is statistically distinct from all other samples (Pr > F = 0.001; Table S23), emphasizing the effect that removal of the dominant fish has on this group.

The correlation matrices of behavior and neural activity data (Figure 4F-I; Table S19) show that the strongest correlations found between all factors are between the different brain areas, emphasizing the interconnectedness of the social decision-making network. Comparing ranks in figure 4F-I, both positive and negative correlations decrease in intensity as rank lowers. Within the social decision-making network, the strongest positive correlations (above 0.75) across ranks are between the PAG and the POA, between the vTn (AH) and the PAG, between the Vv (LS) and the POA, and between the Vv (LS) and the Vs (BNST/meAMY). The aTn correlates negatively with all other social decision-making network nodes, most notably the vTn (AH), Vs (BNST/meAMY) and the PAG, which is particularly evident in rank 2 fish (Table S19). Standout brain-behavior correlations include a strong negative correlation between the PAG and escape behavior, as well as a strong positive correlation between POA activation and rushing behavior.

## Discussion

An area of ongoing investigation in social hierarchy and dominance research is understanding how dominance hierarchies relate to individual physiological and neurobiological states (48). In the current study, behavioral and neural correlates of social hierarchy formation were investigated in a sex-changing fish. We hypothesized that size (i.e., standard length) is the main determinant of rank and dominance, with larger fish being more likely to score higher in both categories. Indeed, dominance hierarchies were rapidly established upon group formation in a size-dependent manner, and each rank was found to have a significantly different dominance score (DS). Fish size significantly affected aggressive and submissive behaviors, and therefore overall DS, confirming the importance of size in dominance hierarchies (15,49). While these findings are not unexpected based on the hierarchy formation in other species (49), and with larger individuals initiating sex change among sequential hermaphrodites (2), our study is one of very few displaying the extent of behavioral rank differences in all group members of a protogynous species (49) and sequential hermaphrodites overall (50–53). This study also differs from others in that we defined rank by behavioral attributes rather than assuming rank based on size.

Surprisingly, ranks 1 and 2 did not differ significantly in aggressive behaviors (Figure 1A). However, this is likely because rank 1 fish spent a large proportion of time in the shelter, displaying how being at the top of a hierarchy allows for priority access to certain resources. This furthermore explains why all other ranks spent significantly more time swimming and on the floor than the dominant individual. It is typical in many social species for the dominant individual to take possession of the shelter, as well as other resources before subordinates (19,54,55). Occupation of the shelter by the dominant fish also corresponds with spotty behavior in the wild (56). In line with this, time spent actively swimming increased as rank decreased, suggesting that locomotion could be a sign of subordination. The energetic cost that comes with swimming versus resting in the shelter further emphasizes the value of being a dominant individual. However, it should be noted that this experiment took place in a laboratory setting with only one shelter being present, potentially making the energy the dominant fish needs to surveil and maintain the territory much less intensive.Although rank 1 and rank 2 fish did not differ significantly in aggression, they did differ in submission. Rank 1 fish showed little to no submission, which is expected in a stable hierarchy without competition and commonly observed in despotic hierarchies, where one individual is dominant over all others (57). Submissive behavior increased as the hierarchy descended but was not always significantly different between individuals. These observations show the importance of taking multiple behaviors into account when studying hierarchy formation. Future studies with spotty wrasse should aim to investigate hierarchy type and which individuals are interacting with each other (i.e., heterarchy) to gain further understanding on their social structure (6). Overall, the current study suggests that spotty wrasse form a linear hierarchy, similar to other protogynous fish (49), where the top individual dominates over all others, the 2^nd^-ranked fish over all but the dominant, and so on. However, further analyses would need to be conducted to confirm linearity (6,58,59).

Disruption of the hierarchy by removing the dominant fish induced significant behavioral changes. Most notably, the rank 2 fish displayed significantly increased aggressive behavior (Figure 2), as well as a significant decrease (in some cases, complete absence) of submissive behavior. This type of behavior has been investigated in many other sex changing species, with the removal of the dominant fish causing rapid changes in behavior (9,60,61). Most fish displayed increased activity over this period, with all ranks decreasing time resting on the floor and some a significant increase in swimming, demonstrating the disruption that has occurred. In the control treatment, where the dominant fish was removed and immediately returned, fish did not seem to have changes in their state behavior except for rank 1 (the displaced fish). Rank 1 showed a significant increase in its time swimming and decrease of time spent in the shelter. Rank 1 also showed an increase in its aggression with a significant increase in rushing behavior. This could be because rank 1 is trying to reaffirm its dominance and the stress caused by its removal. Behavior of rank 1 likely explains why some of the other fish experience a decrease in DS or increase in submissive behavior as well.

All animals exhibit adaptive responses to their social surroundings through a complex neural framework (31), primarily being: 1) the social behavior network (62,63), and 2) the mesolimbic reward system (64,65). The differentially activated brain areas in this study are all part of the mesolimbic reward system (Vv, Vs, Vd, TPp, Vc and Dm) and/or the social behavior network (aTn, vTn, PAG, POA, Vv and Vs) (Figure 3, Table 17c). These regions (except for aTn) all showed increased activation in the rank 2 fish compared to all other fish, after social disruption (Figure 3). Further, we also show that that there is higher positive correlation between aggressive behaviors and the social behavior network in rank 2 fish compared to the lower ranked fish (Figure 4). These findings support previous research that the activation of the social behavior network occurs during social ascension in vertebrates(66) and provide further evidence for the role of the social decision-making network (SDMN) in the initiation of reproductive behavior changes. The social behavior network is generally believed to regulate behaviors like reproduction, aggression, and parental care through interactions with various sex steroid and neuropeptide hormones (62,63), whereas the mesolimbic reward system assesses the salience of stimuli through dopaminergic signaling (64,65). The current study ties in well with previous studies on the initiation of socially-controlled sex change, as well as hierarchy formation in general (67,68). A multitude of neuroendocrine factors have been shown to be necessary, but not sufficient for sex change, including GnRH (69), arginine-vasotocin (70), cortisol (71), and sex steroids (52,72,73). Our data suggest that the onset of socially-controlled sex change does not depend on one specific endocrine factor, produced in one specific brain area, but rather involves a highly interconnected network of brain areas and their associated signaling factors. Although pS6-immunohistochemstry is a reliable readout of neural activity, it does not distinguish cell type. We suggest that future work should focus on the transcriptional profile of specific cell populations within the social decision-making network.

The one brain region with a specific decrease in neural activation upon attaining dominance (i.e., in rank 2 fish vs all other fish) was the aTn, which was negatively correlated with other regions of the SDMN (Figure 4C-F and Table S19). The mammalian homolog of the aTn, the ventromedial nucleus of the hypothalamus, drives the switch between feeding and defensive behaviors (74). The decreased activity in this area could therefore be an indication of active suppression of unhelpful behaviors during attainment of dominance. Interestingly, a recent study reported an aTn-specific difference in pS6 immunoreactivity between male and female cichlids in an assay of aggressive behavior (75). In accordance, Luong et al. (30) found that in the closely related bluehead wrasse (*Thallasoma bifasciatum*) stimulation of brain areas including the Vs, Vv, Vd and POA induced changes in coloration associated with courtship behavior. In the same study pS6 and c-fos immunohistochemistry revealed that the interpeduncular nucleus, red nucleus, and ventrolateral thalamus were more active in subordinate (i.e., all but the TP and dominant IP) fish. These regions were not found to be significantly more active in subordinate fish in our study; however, Luong et al. did not differentiate subordinates into ranks. The difference is likely due to the differential brain sampling timeline used: whereas non-courting animals in the Luong et al. study were observed for at least four hours before sampling, we euthanized our fish 1 hour after social disruption.

In conclusion, our work shows that New Zealand spotty wrasse rapidly form social hierarchies, based mainly on size. Disruption of this hierarchy induces rapid (1hr) and substantial changes in behavior and leads to differential activation of the social decision-making network in the brain. This study confirms the importance of size in social group dynamics and provides the behavioral and neuroanatomical base for further investigation of social hierarchy formation, as well as the initiation of socially-controlled sex change.

## Author Contributions

Experimental design was constructed by H.Q., C.v.d.B., K.K, N.G., C.B., S.M., E.T. and C.J. Behavioral experiments were carried out by H.Q., C.v.d.B, K.K. and S.M. Behavioral data were watched and generated by H.Q. Brain region data were generated by K.K. Brain and behavior data were analyzed by H.Q., C.v.d.B and K.K., with statistical and methodological input from C.B. and N.G. Figures were generated by H.Q., C.v.d.B. and K.K. The manuscript was written by H.Q., C.v.d.B and K.K. and edited and approved by all authors. H.Q., C.v.d.B. and K.K share co-first authorship.

## Acknowledgments

The authors would like to acknowledge the assistance and technical support of Kevin Green, Tessa Hamer, and Ethan Russell.

## Data Availability Statement

The consolidated data for the behavioral datasets generated for this study can be found in Supplementary Table S2 (Group formation) and Table S3 (Social disruption). Raw data files from BORIS can be found at https://figshare.com/s/54488470957b773b34f5.

