## Supplementary material for "Behavioral and neural correlates of social hierarchy formation in a sex-changing fish"

**Contents**

**Supplementary Figure S1.** Individual mass and average standard length (SL)

**Supplementary Figure S2.** Diagram of Experimental Design.

**Supplementary Figure S3.** pS6 antibody specificity.

**Supplementary Figure S4**. Counterstaining of ROIs

**Supplementary Figure S5.** Dominance scores by average standard length

**Supplementary Figure S6.** Average standard length by rank

**Supplementary Figure S7.** Dominance scores by each individual observation (chronological).

**Supplementary Figure S8.** Gonadal histology.

**Supplementary Figure S9**. Scree plots of principal components vs eigenvalue for behavior and neural data post social disruption.

**Supplementary Figure S10**. Comparative principal components of neural and behavioral factors post social disruption.

**Supplementary Materials Notes**

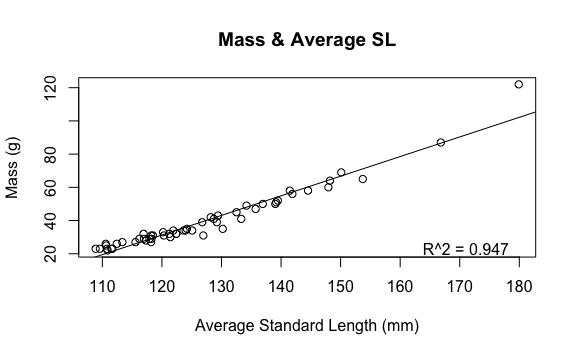

**Supplementary Figure S1.** Individual mass and average standard length (SL).

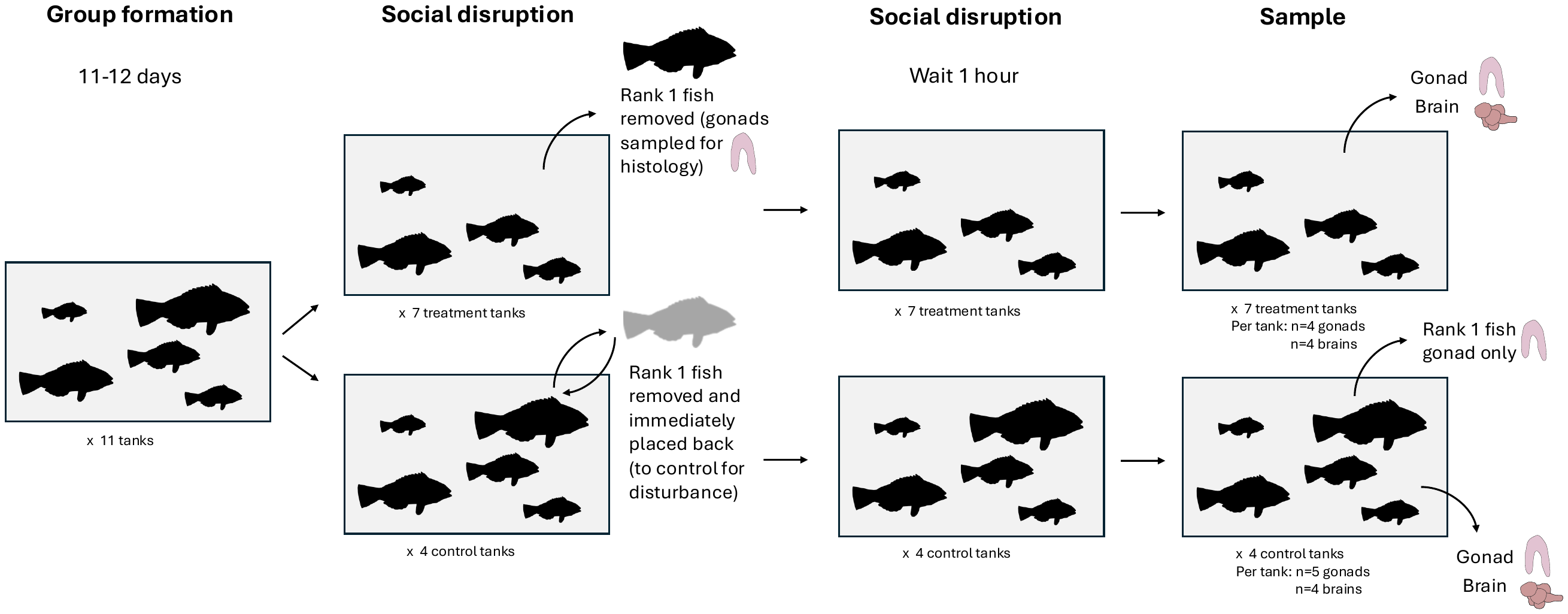

**Supplementary Figure S2.** Diagram of Experimental Design.

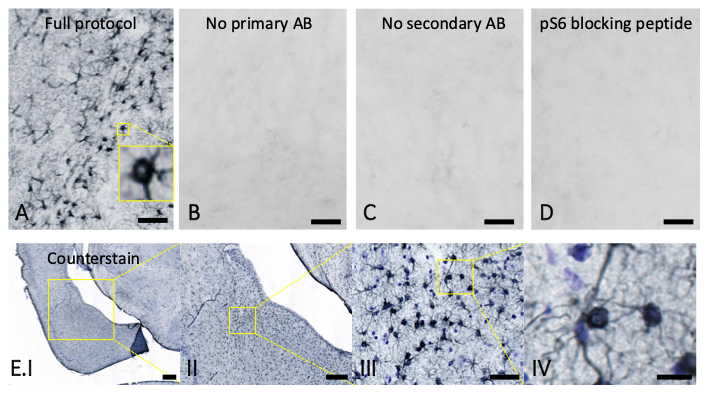

**Supplementary Figure S3.** pS6 antibody specificity. pS6 staining produces clear distinctly stained cells (A) but staining is absent when primary (B) or secondary (C) antibody is left out. Staining is completely eliminated when the antibody is pre-absorbed with a pS6 blocking peptide (Cell Signalling #1220) (D). A-D: Scale bar = 20µm. Nissl counterstaining (in blue) showing overlap between Nissl-stained cells and pS6 stained cells (in black), as well as Nissl-stained cells which are not also stained with pS6 (i.e., cells are not showing recent neural activation) (E.I-IV). Scale bars of E panels are 100µm (E.I), 50µm (E.II), 20µm (E.III) and 4µm (E.IV).

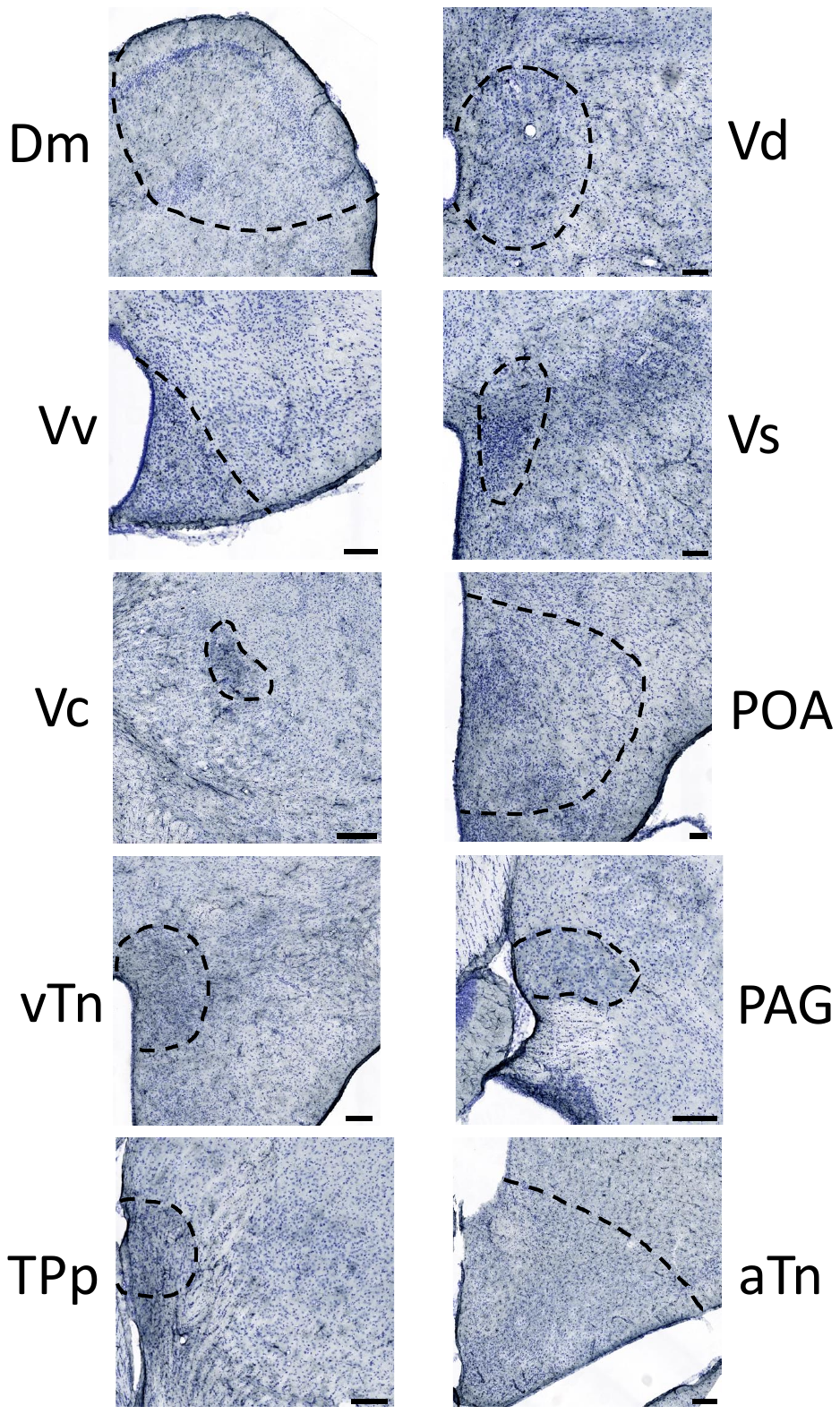
**Supplementary Figure S4**. Counterstaining of ROIs

Brain regions analyzed were identified based on the spotty wrasse brain atlas (Kamstra et al., 2023) and confirmed through Nissl counterstaining. Panels show Nissl substance in blue and pS6 in black. Scale bar = 100µm.

Anatomical location of ROI’s:

Medial part of the dorsal telencephalon (Dm): dorsally in telecephalon, adjacent to dorsal ependymal lining and telencephalic ventricle

Dorsal part of the ventral telencephalon (Vd): medial of central zone of the dorsal telencephalic area

Ventral part of the ventral telencephalon (Vv): medial of medial olfactory tract

Supracommissural nucleus of the ventral telencephalon (Vs): dorsal of commissural anterior, pars dorsalis

Central nucleus of the telencephalic area (Vc): relatively small nucleus situated centrally in telencephalon

Ventral tuberal region of the hypothalamus (vTn): less ventral than suggested by it’s name, adjacent to the diencephalic ventricular space

Preoptic area (POA): adjacent to the diencephalic ventricular space, posterior of hypothalamic area

Periaqueductal gray (PAG): ventral of superior colliculus in midbrain

Periventricular nucleus of the posterior tuberculum (TPp): dorsal of ventral hypothalamic area

Anterior tuberal nucleus (aTn): ventrally in central hypothalamic area, ventral of central posterior thalamic nucleus

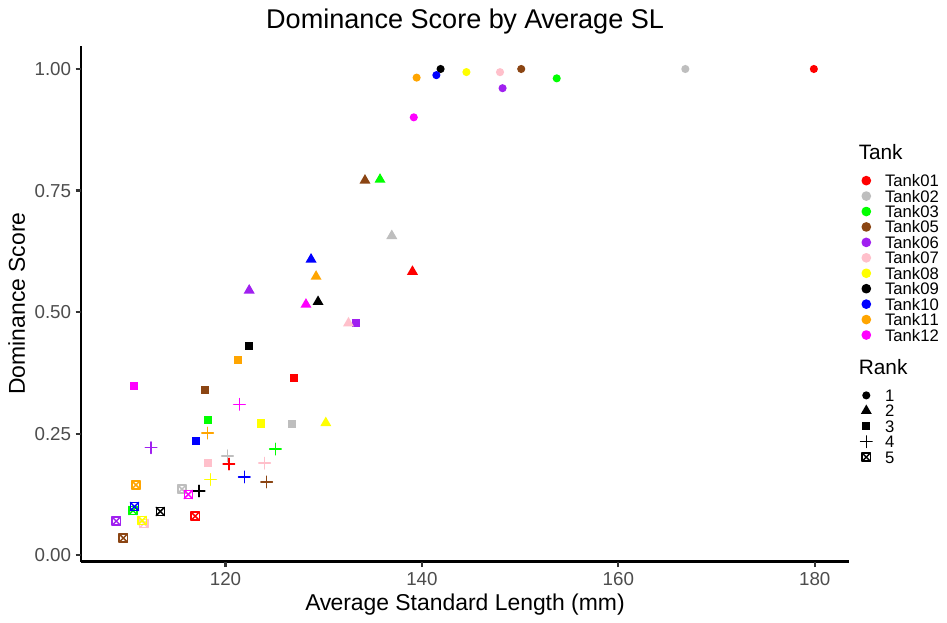

**Supplementary Figure S5.** Dominance scores by average standard length. The average SL has a positive relationship with the dominance score. Each point represents an individual fish, with color signifying which tank each was in and the shape signifying the rank it held.

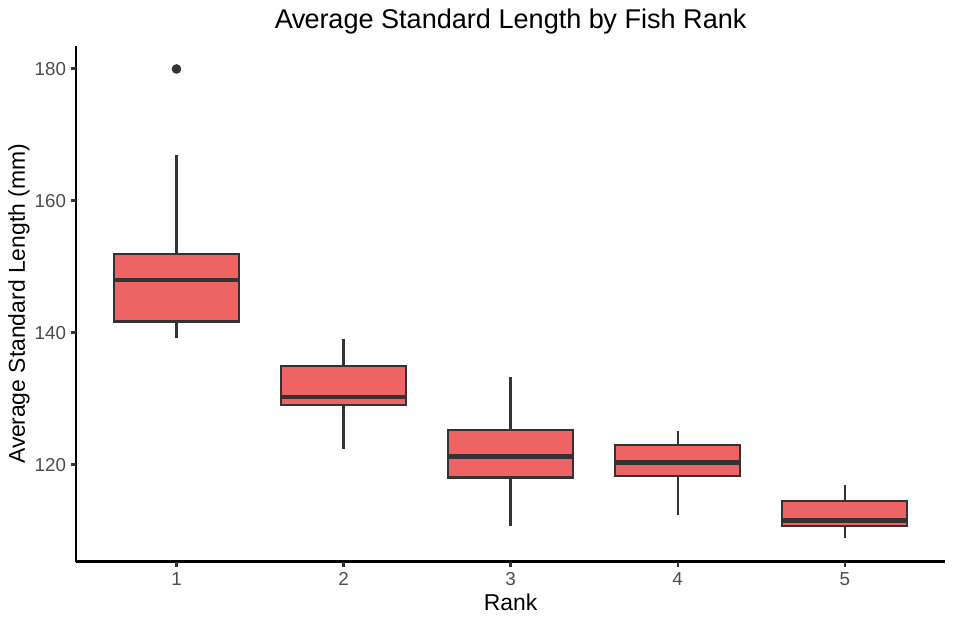

**Supplementary Figure S6.** Average standard length by rank. Each rank differs in average SL from each other, except for ranks 3 and 4.

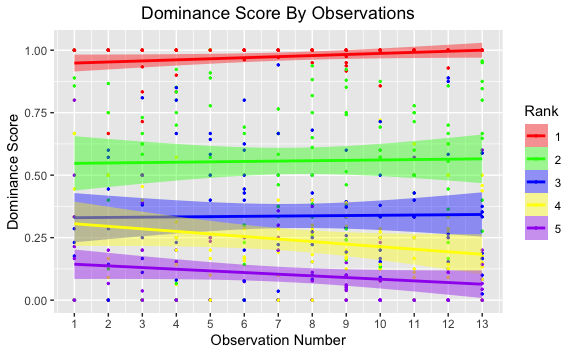

**Supplementary Figure S7.** Dominance scores by each individual observation (chronological).

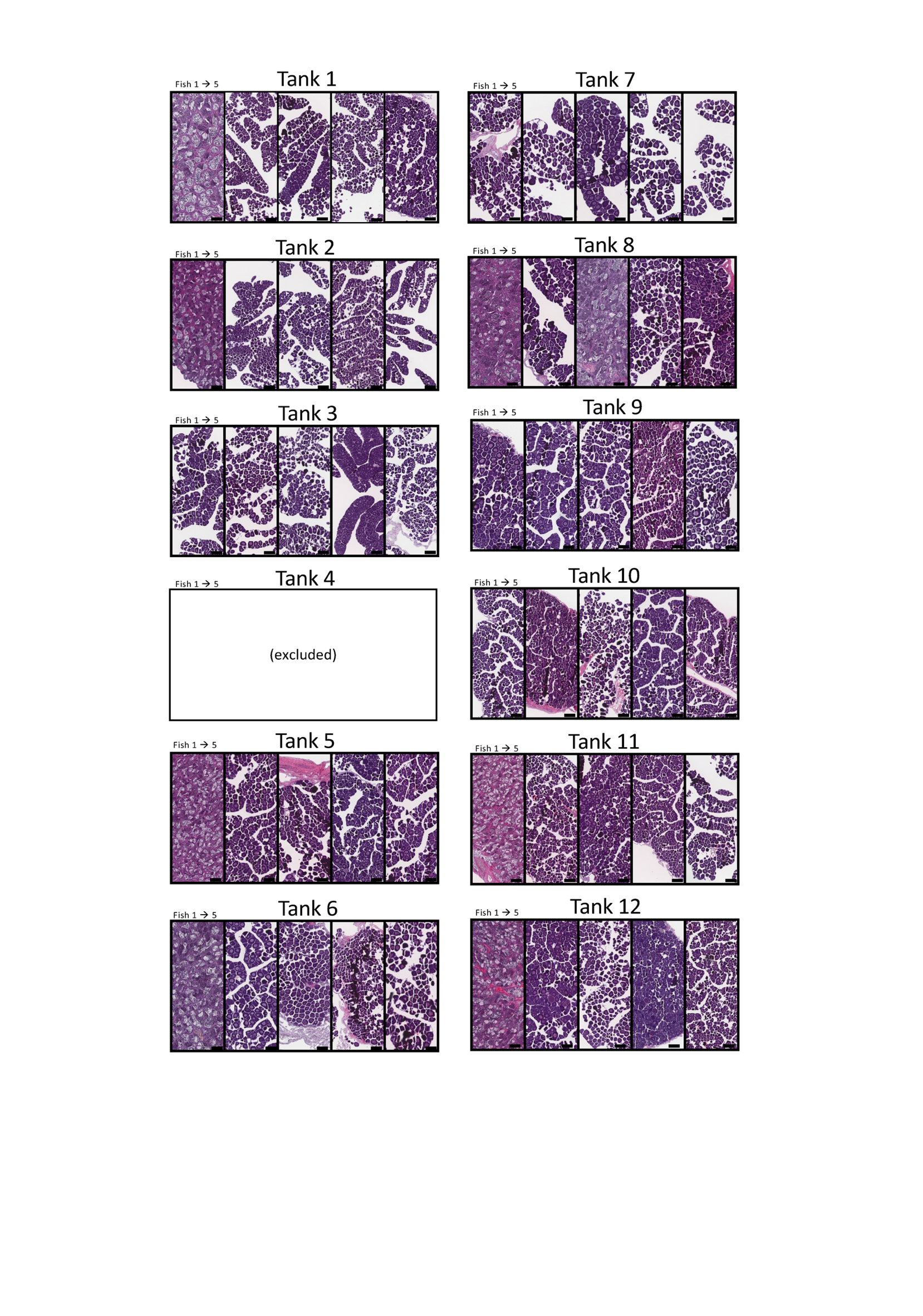

**Supplementary Figure S8.** Gonadal histology. Fish are ordered left to right by rank. Tank 4 was excluded entirely from analysis due to technical reasons.

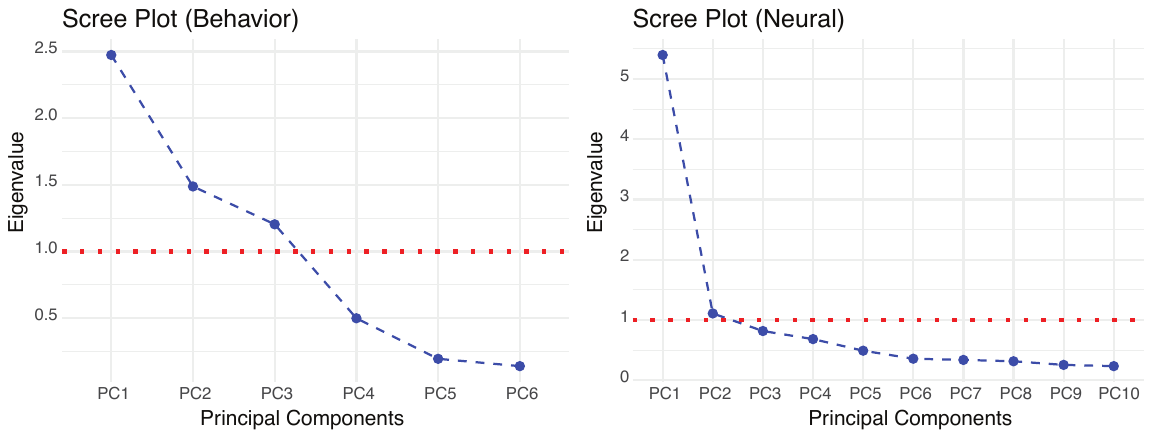
**Supplementary Figure S9**. Scree plots of principal components vs eigenvalue for behavior and neural data post social disruption. Principal components with an eigenvalue >1 were further analyzed (see plots below).

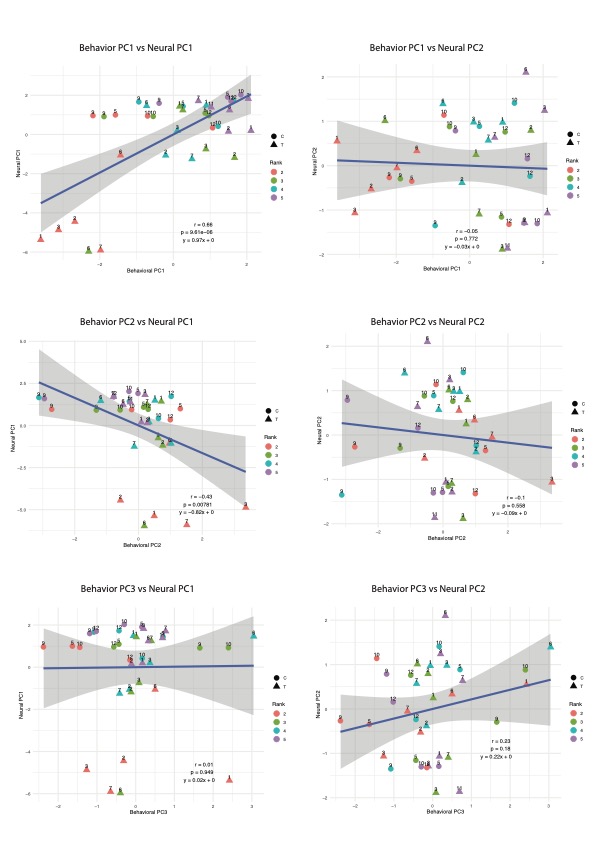
**Supplementary Figure S10**. Comparative principal components of neural and behavioral factors post social disruption. Sample points are labelled by tank. C= control, T= treatment. Only PCs with an eigenvalue > 1 were considered (see scree plots above). Behavioral PC1 vs Neural PC1 is the only comparison to show a significant positive linear relationship. Statistical significance calculated using Pearson's product-moment correlation.

**Supplementary Materials Notes**

**Note on gonad histology results**: Hierarchical status was indicated not only by behavior and neural activation patterns, but also by gonadal histology. After only 12 days of group formation, 7/11 of the dominant fish were found to have clearly masculinized gonads (Figure S8). A challenge working with New Zealand spotty wrasse is the potential to inadvertently introduce female-looking IP males in your experimental groups. Theoretically, the 7/11 fish that showed masculinized gonads in Figure S8 above, while also being the biggest and most dominant fish in their tank, could all be IP males. However, since the distribution of IP males in natural populations (~10% of fish) is random (e.g. not size-skewed), it would be expected that any IP males in our tanks would be distributed randomly as well. It is statistically not unlikely that 8 out of 60 fish are IP male, but the statistical likelihood of 7 out of those 8 fish being the biggest and most dominant fish in their tank is 0.032%.

The most likely explanation for the masculinization of these gonads is that that larger IP fish had already begun gonad masculinization before groups were initiated. Given the short time frame, rapid sex change is a less likely explanation, and the likelihood of all of the largest fish being inadvertently introduced IP males is also statistically low. The behavior of these fish during hierarchy formation also did not differ from other feminized dominant fish and there was no statistical difference in fish size between masculinized fish and feminized fish. However, if differences in their behavior during social disruption exist there is insufficient statistical power to test this, with only two of the non-masculinized fish in the treatment condition and two in the control condition for social disruption. The potential for social behavioral differences between groups with dominant fish with and without masculinized gonads, including how the dominant fish communicates dominance and what neuroanatomical or genetic signatures of dominance exist, is worthy of further investigation, but outside the scope/power of this manuscript to test.

**Note on “outlier/confounding” behaviors for Figure 4 (brain-behavior correlation analyses):** During the course of the Group Formation and Social Disruption filming sessions, many different behaviors were observed and analyzed in BORIS, however, not all were relevant for use in the correlation analyses. For example, the behavior “Out of view” of the camera was used to adjust other behaviors to a proportional time, and is therefore not relevant as a behavior on its own. The “feeding” behavior was also excluded as this was only measured during Group Formation, not Social Disruption. Other behaviors, such as “Head bob” and “Chase follow” had too few observations in the Social Disruption experiment (4/37 fish and 2/37 fish, respectively) to include in subsequent analyses. Lastly, some behaviors were confounding and were therefore excluded from further analysis - for example, we initially measured “attack” as an individual behavior, which was essentially defined as a rush where the attacking fish made contact with another fish. However, this behavior was not observed as frequently as rush behaviors, whereby the attacking fish ‘rushed’ at another fish but did not necessarily make physical contact. Therefore, we decided to continue analyses using the rate or ‘amount’ of rush behavior exhibited by a fish, but not the intensity of that behavior (e.g., whether the rush resulted in contact).

**Note on tank 8**: In tank 8, it was found that two fish had masculinized gonads (Rank 1 and Rank 3 by size; Figure S8), leading us to believe that the 3rd-ranked fish in this tank was an IP male. While there is insufficient data here to statistically determine if IP males behave differently to IP females in the context of hierarchy formation, anecdotally, the behavior we observed of the IP male was more aggressive than was expected for its rank based on size. Tank 8 was therefore excluded from further analyses.
